# Zyxin and non-muscle myosin are required for single fibroblast durotaxis, but Rho-kinase activity and the Arp2/3 complex are dispensable

**DOI:** 10.1101/2022.06.03.494725

**Authors:** Reem M. Hakeem, Bhagawat C. Subramanian, Max A. Hockenberry, Zayna T. King, Mitchell T. Butler, Wesley R. Legant, James E. Bear

## Abstract

Durotaxis, migration of cells directed by stiffness gradient, is critical in development and disease. To study the molecular determinants of single cell durotaxis, we developed an all-in-one photopolymerized hydrogel system containing areas of stiffness gradients with different slopes, along with uniform stiffness (soft and stiff) regions. We find that fibroblasts rely on non-muscle myosin II (NMII) activity and the LIM-domain protein zyxin for durotaxis on both steep and shallow stiffness gradients. Importantly, unlike haptotaxis, the Arp2/3 complex is dispensable for durotaxis on both stiffness gradients. Lack of Arp2/3 results in a filopodia-based durotactic migration that is equally efficient as that of lamellipodia-based durotactic migration. Finally, we reveal an essential role for the actin-bundler fascin in the formation and asymmetric distribution of filopodia during filopodia-based durotaxis in shallow, but not steep, stiffness gradient. Together, our all-in-one hydrogel system can serve as a platform to identify, discriminate, and characterize stiffness gradient specific molecular mechanisms that cells employ to efficiently durotax.

## INTRODUCTION

Cells actively probe and respond to mechanical cues in their microenvironment (van Helvert et al., 2018). The term ‘durotaxis’ was first coined to describe the ability of fibroblasts to preferentially migrate towards the stiffer side of biphasic stiffness polyacrylamide hydrogels (Lo et al., 2000). Since then, studies have demonstrated that stiffness-directed migration of cells is essential during development and morphogenesis (Thompson et al., 2019; Shellard and Mayor, 2021), immune response (Bollmann et al., 2015), fibrosis (Liu et al., 2010), and cancer progression (McKenzie et al., 2018; DuChez et al., 2019), where cells migrate through diverse mechanical environments. Several cellular mechanisms and molecular pathways have been implicated in mechano-sensing during durotaxis such as non-muscle myosin II (NMII) activity (Raab et al., 2012; Sunyer et al., 2016), fluctuating focal adhesions (FAs)(Plotnikov et al., 2012), filopodia-based extensions (Wong et al., 2014), Arp2/3 dependent-lamellipodia (Oakes et al., 2018), asymmetric focal adhesion kinase activity (Lachowski et al., 2018), and the Golgi-derived microtubule regulation of FAs (Rong et al., 2021). However, the ways in which cells effectively integrate such processes and molecular pathways to sense, polarize and directionally navigate along stiffness gradients remains poorly understood. (SenGupta et al., 2021)

One impediment to a fuller understanding of durotaxis is the lack of simple and robust experimental approaches to generate stiffness gradients. While several approaches have been developed, they are limited by technical and conceptual shortcomings that hinder progress. The simplest and oldest assay uses two droplets of polymerizing acrylamide solutions (with different stiffness properties), brought together in coverslip sandwich to produce a binary stiffness gradient (Lo et al., 2000; Raab et al., 2012; Lachowski et al., 2018). However, this system produces a very short gradient region and lacks precise tunability. With the introduction of photopolymerized polyacrylamide gels, more precise control became possible, but the implementation of this approach thus far has involved mechanically moving masks that must be precisely controlled in a non-linear fashion or fixed neutral density filters that limits gradient properties (Sunyer et al., 2016; Evans et al., 2018; DuChez et al., 2019; Rong et al., 2021; Yip et al., 2021). Another approach uses physical stretching or deformation of polyacrylamide gel with glass pipettes attached to micromanipulators near individual cells to induce short distance cell migration (Plotnikov et al., 2012; Svec et al., 2019; Puleo et al., 2019). However, it is not clear what kind of stiffness gradient this approach generates as these have not been measured. Mechanical pulling on the substrate also applies extrinsic forces on nearby cells, leading to events such as opening of stretch-activated ion channels, which may be quite different from exerting their own intrinsic pulling/pushing forces on substrates of varying stiffness, leading to durotactic sensing and directed migration.

To address this knowledge gap, we developed a robust experimental system to produce stiffness gradients, allowing us to test the roles of key molecular players and cellular processes in single cell durotactic migration. This system is simple to implement and produces both steep and shallow gradients that capture a physiologically relevant substrate stiffness range (2-75 kPa) (Irianto et al., 2016) in a single engineered hydrogel system. In addition to gradients of stiffness, these hydrogels also contain areas of uniform stiffness (both stiff and soft) that serve as internal controls for the durotaxis responses observed in the gradient regions. Finally, this system is amenable to both live and fixed cell fluorescence imaging on a standard confocal microscope. Using this system, we explored the basic requirements of single cell durotaxis in fibroblasts and revealed critical molecular regulators of durotaxis. While several processes and components such as NMII activity and the mechano-sensing protein zyxin are required for durotaxis, other components such as Rho-kinase activity and the Arp2/3 complex are surprisingly dispensable for this process. Together, our work explores the critical molecular mechanisms underlying durotaxis in single cells and lays the foundation for future investigations of this process.

## RESULTS

### Development of a photopolymerization-based durotaxis assay

To study single cell durotactic migration, we developed a simple and highly reproducible photopolymerizable hydrogel assay with tunable gradients of stiffness that captures the range of stiffnesses observed in mammalian tissues (2-75 kPa) (Irianto et al., 2016) (**Figs. 1A and S1A**). We used lithium phenyl-2,4,5-trimethylbenzoylphosphinate (LAP), a photoinitiator that cleaves into substituent radicals upon exposure to 365 nm light, to initiate polymerization of polyacrylamide hydrogels (Fairbanks et al., 2009; Purbrick, 1996). The LAP photo-initiator provides spatial and temporal control of polymerization to generate stiffness gradient acrylamide gels (Fairbanks et al., 2009). Briefly, a drop of acrylamide/bis-acrylamide/LAP mixture was sandwiched between two coverslips and positioned with a 3D-printed holder beneath a house-shaped mask. The asymmetric house shape allowed for rotational orientation of the translucent gradient gel during subsequent imaging. The lower half of the house shape was masked (**Fig. 1A**), and the upper half was exposed to 365 nm light for 3 min to establish a uniform stiff area in the ‘roof’ region of the gel. After 3 mins of exposure, the temporary mask was removed, and the entire house shape was exposed for an additional 45 secs to allow polymerization of the lower region of the gel. During illumination of the upper half of the house shape, a steep gradient at the boundary of illumination was formed, likely through the diffusion of activated LAP from the illuminated region of the gel sandwich into the non-illuminated area. At the bottom of the house-shaped gel, where the mixture is only illuminated for 45 secs, a uniform soft area formed (**Fig. 1A**). By differential interference contrast (DIC) microscopy, we observed a distinct line of altered contrast that appeared to correspond to the boundary of initial illumination (**Fig. 1B**). This line of altered contrast fortuitously served as a fiduciary point of reference with respect to the gradient generated in this system during subsequent imaging experiments.

**Figure 1.**
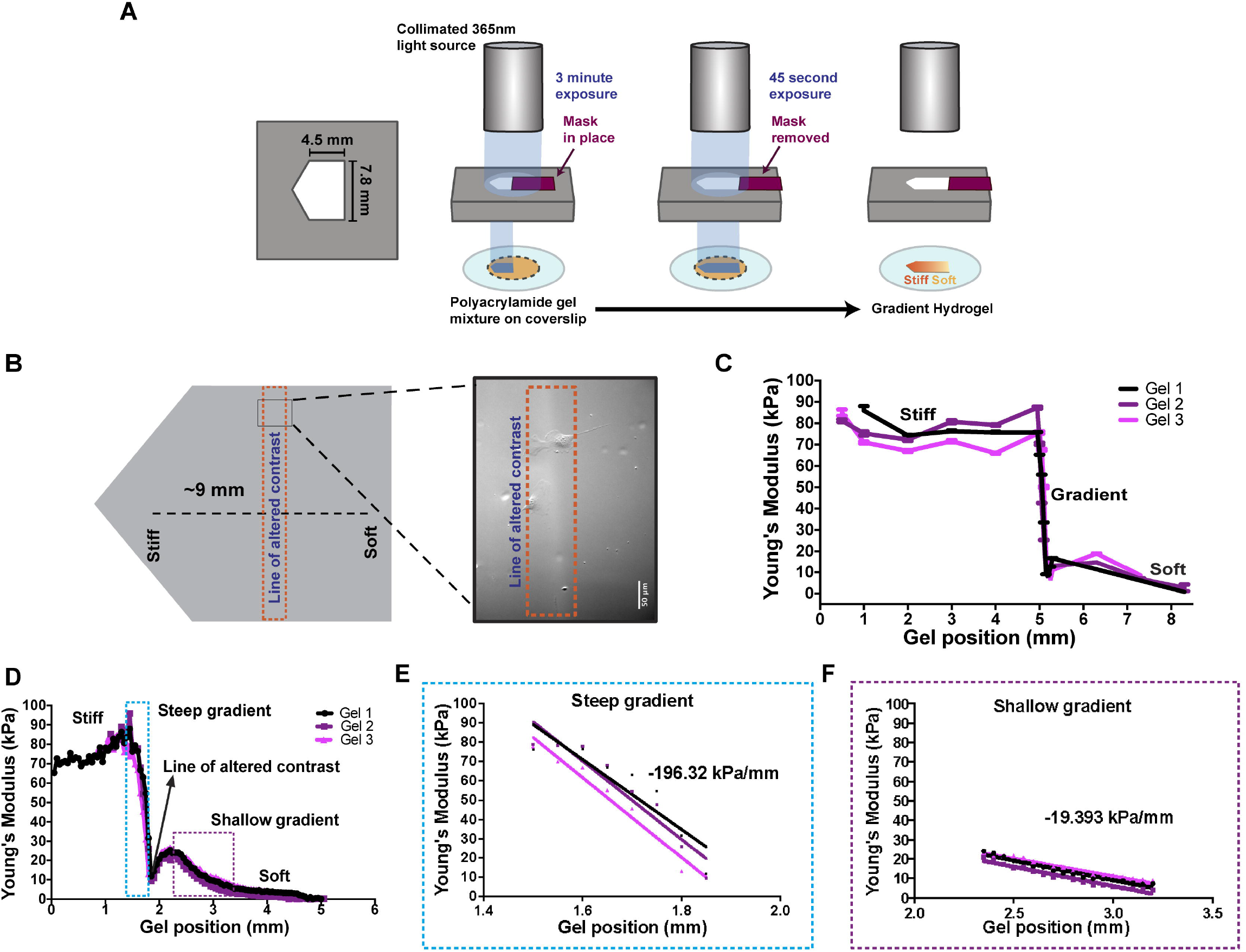
Generation of a novel photoinitiation-based hydrogel system to study single cell durotaxis. **(A)** Schematic representation of the various steps involved in the generation of polyacrylamide-based house-shaped hydrogels using the photoinitiation approach. **(B)** Pictorial depiction of ‘the line of altered contrast’ (dashed orange rectangle) with respect to the length (dashed black line) of house-shaped hydrogel encompassing stiff and soft regions (on the left). Region within the dashed black rectangle is highlighted in a representative DIC image revealing ‘the line of altered contrast’ at the center of hydrogel (on the right). **(C)** AFM-based measurements of the elastic modulus distribution across the length of the engineered hydrogel, as determined by the Hertzian Contact Model, is plotted. All error bars are s.e.m. (n=3 independent hydrogels). **(D)** Elastic modulus distribution of the hydrogel as obtained via nano-indentation method (n=3 independent hydrogels) is plotted. Different identified regions are indicated, dashed rectangular cyan and magenta box highlight the steep and shallow stiffness gradients respectively and the arrow points to the line of altered contrast. **(E-F)** Linear regression analysis of the values obtained from three independent hydrogels using nanoindentation is plotted and fitted to line for steep stiffness gradient (E) and shallow stiffness gradient (F). Each line represents a line of best fit for each individual hydrogel. The value indicated is the pooled slope of 3 hydrogels.

We characterized the elastic modulus profile of the generated house-shaped hydrogel using atomic force microscopy (AFM). As expected, the gel displayed a uniform stiff region (~75kPa) at the top, a steep gradient at the center (~75-10kPa), and a uniform soft region (~2kPa) at the bottom (**Fig. 1C**). Interestingly, we observed a more complex pattern of stiffness in the area below the steep gradient that included an area with a secondary gradient of stiffness that was much shallower. Since AFM did not allow us to simultaneously visualize the gel and make stiffness measurements, and was limited in its lateral spatial resolution, we resorted to using a nano-indentation approach for a more precise measurement of stiffness across the gel. Using this approach, we characterized the stiffness profile of the house-shaped hydrogels polymerized onto 500 μm^2^ gridded coverslips with 50 μm lateral resolution (**Fig. S1B**). Using the grid lines for reference, we obtained a correlative elastic modulus profile of the gel, relative to the line of altered contrast visualized by light microscopy (**Fig. 1D**). This approach revealed that the line of altered contrast was a local minima of stiffness and that the hydrogel contained two distinct gradients: a steep gradient (196 kPa/mm) ranging from 10 to 75 kPa, ~400-500 μm in length directly above the line of altered contrast **(Figs. 1D-E)**, and a shallow gradient (19 kPa/mm) that ranged from 5 to 20 kPa, over 850 μm in length and beginning ~500 μm below the line of contrast (**Figs. 1D and 1F**). This profile of stiffness was highly reproducible and allowed us to generate two-gradient hydrogels with areas of uniform stiff and soft regions serving as internal controls in each hydrogel.

### Fibroblasts display single cell durotaxis on stiffness gradients

To assess single cell durotaxis using this hydrogel system, we used JR20 dermal fibroblasts, a diploid subclone derived from the previously published bulk population (Rotty et al., 2017), seeded on fibronectin-coated hydrogels. To ensure that fibronectin coating on the hydrogels was even and not influenced by the stiffness gradients, we first analyzed the intensity profile of Cy5-conjugated fibronectin across the house-shaped gel and found a uniform distribution of fibronectin across the hydrogel **(Figs. S1C-D)**. Cells were then seeded on the fibronectin-coated gels and imaged by DIC microscopy for up to 24hrs **(Fig. 2A)**. From the computed single cell tracks, we calculated the standard parameters of velocity and persistence (D/T), as well as forward migration index (FMI; Cos(θ)*D/T) (Wu et al., 2012), a measure of directionality that is independent of cell velocity. Cells displayed comparable velocities across various regions of the hydrogel (**Fig. 2B**), with an elevated persistence on the steep gradient region (**Fig. 2C**). On both the shallow and steep gradient regions, cells displayed a consistent directed migration up the gradient that is reflected in the positive average FMI values whose 95% confidence intervals did not contain zero (**Fig. 2D**). To our surprise, both gradients provoked a similar degree of directed migration (average FMI), suggesting that fibroblasts, migrating as single cells, are highly responsive to gradients of substrate stiffness across a range of stiffnesses and slopes.

**Figure 2.**
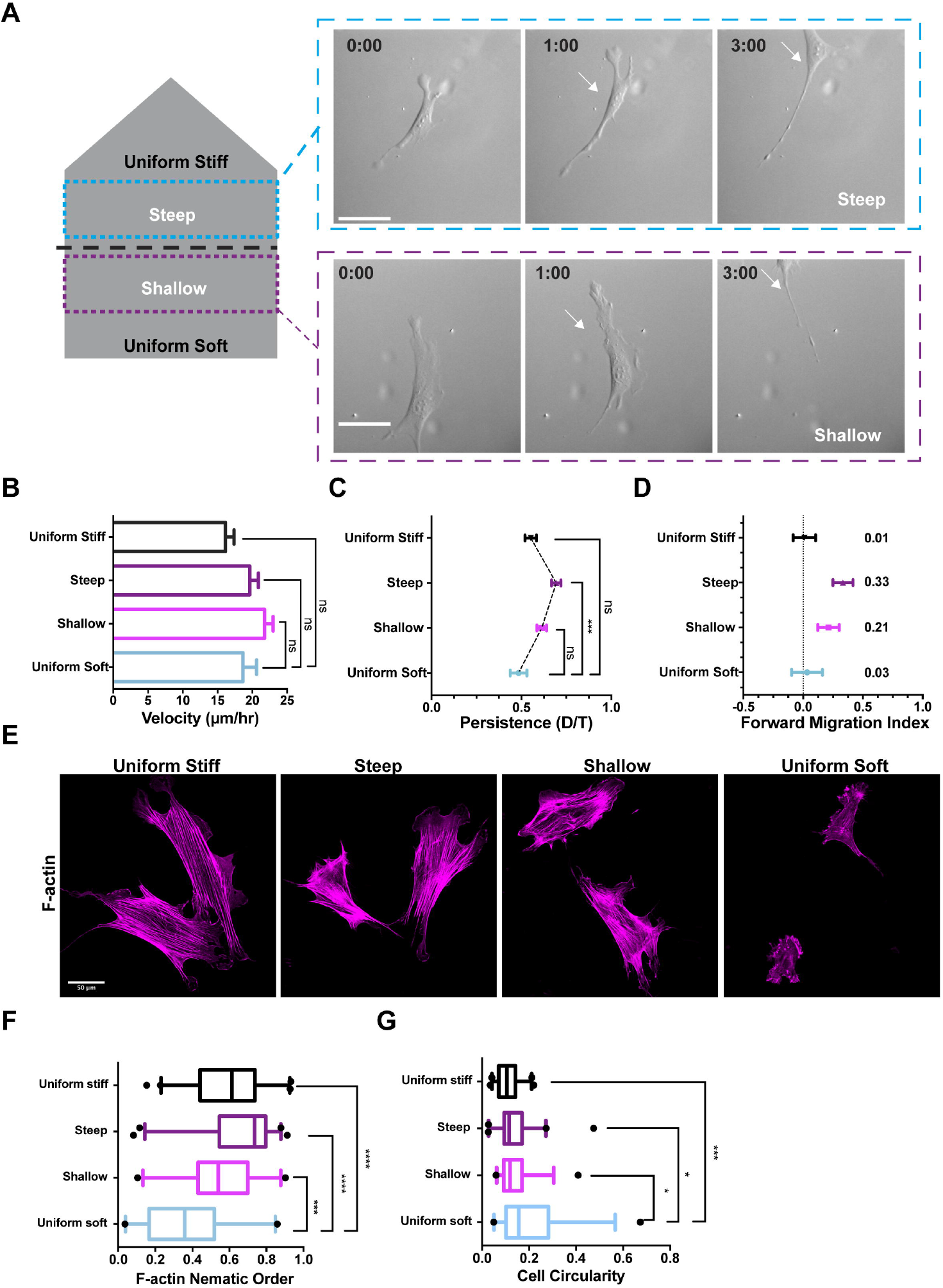
JR20 dermal fibroblasts display single cell durotaxis with polarized Factin alignment on stiffness gradients, but not uniform stiffness regions. **(A)** Left: pictorial illustration of the identified uniform stiff, steep gradient, shallow gradient and uniform soft regions in the engineered hydrogel system. Steep and shallow stiffness gradient regions are highlighted using cyan and purple dashed rectangles respectively. Right: Representative timelapse DIC images of JR20s migrating in the steep (top row) and shallow (bottom row) stiffness gradients. Time stamp in h:min and 0:00 marks the time when cells initiated migration during the acquisition. White arrow indicates the relative position of cells with respect to field-of-view (FOV); Scale bar, 50 μm **(B-D)** Quantification of the velocity (B), persistence (C) and FMI (D) of JR20s in various regions of the hydrogel is plotted. Values obtained from six independent experiments are plotted with ‘n’ representing individual cell value in each region - uniform stiff (n=82), steep (n=105), shallow (106) and soft (n=36) regions. Velocity and persistence are plotted as mean ± s.e.m. FMI is presented as values ± 95% confidence interval. Significance was determined using One-way ANOVA with Tukey’s multiple comparison for velocity and Persistence. **(E)** Representative confocal images of phalloidin-stained JR20s from different regions of the hydrogel. Scale bar, 50 μm. **(F)** Quantification of the F-actin nematic order for individual phalloidin-stained JR20s from 4 independent experiments with ‘n’ representing individual cell value in each region - uniform stiff (n=40), steep (n=44), shallow (36) and soft (n=28) regions is plotted. Values are plotted as 5th-95^th^ percentile of mean and one-way ANOVA with Tukey’s multiple comparison test was performed to determine statistical significance. **(G)** Analysis of the cell circularity across uniform stiff, steep, shallow, and uniform soft regions of the hydrogel. Data represents values from 4 independent experiments, with ‘n’ representing individual cell value in each region - uniform stiff (n=40), steep (n=42), shallow (36) and uniform soft (n=28) regions. Data points in box and whiskers plot represent the 5th-95^th^ percentile range and one-way ANOVA with Tukey’s multiple comparison test was used to determine statistical significance.

One of the advantages of our all-in-one hydrogel system is that we can image molecular components within cells and compare them across different stiffness regimes or gradients within the same experiment. Since the actin cytoskeleton is central to cell shape and motility, we first stained for F-actin (**Fig. 2E**). Actin stress fibers appeared well aligned on uniform stiff and the steep gradient region and progressively less well organized on the shallow gradient followed by uniform soft regions. To quantify this effect, we measured the alignment of F-actin structures within cells using the nematic order parameter in the various regions. Consistent with our visual impressions, the F-actin nematic order value increased with the increase in gel stiffness (**Fig. 2F**), in line with previous reports of fibroblasts plated on uniform soft and stiff substrates (Doss et al., 2020). This altered organization of F-actin in different regions also correlated with changes in cell shape that we quantified by measuring circularity, which showed a gradual decrease as the stiffness of the gel increased (**Fig. 2G**).

### Durotaxis relies on non-muscle myosin II (NMII), but not Rho-kinase activity

Using our hydrogel system, we set out to define the molecular determinants of durotaxis using a combination of pharmacological and genetic approaches. Based on the F-actin organization of cells on durotactic gradients and previously published work (Raab et al., 2012; Sunyer et al., 2016), we postulated that actomyosin contractility would be essential for durotaxis. Fibroblasts express two isoforms, NMIIA and NMIIB, that have been shown to cooperatively polarize the actin cytoskeleton to promote migration of mesenchymal stem cells on soft substrates (Raab et al., 2012). To study the role of actomyosin contractile machinery during durotaxis, we inhibited NMII activity using para-amino Blebbistatin (hereafter abbreviated Bleb)(Straight et al., 2003; Várkuti et al., 2016). Consistent with the known role of NMII activity in the maturation of focal adhesion (FA) structures (Kuo et al., 2011; Burridge and Chrzanowska-Wodnicka, 1996), treatment of cells with Bleb, but not DMSO, dramatically reduced vinculin-positive FAs and F-actin structures at the leading edge of cells plated on stiffness gradients (**Fig. 3A**). Treatment with Bleb also resulted in reduced cell velocity and was accompanied by a complete loss in directionality (FMI) on both the stiffness gradients. This effect on velocity and directionality was reversed upon removal of Bleb (wash-out) from the media (**Figs. 3B-C**). To confirm our observations, we also tracked cells before and after addition of Bleb (wash-in treatment) and found a similar impact on FMI on both the gradients upon Bleb treatment (**Fig. S1E**). These data demonstrate that NMII activity is indispensable for durotaxis.

**Figure 3.**
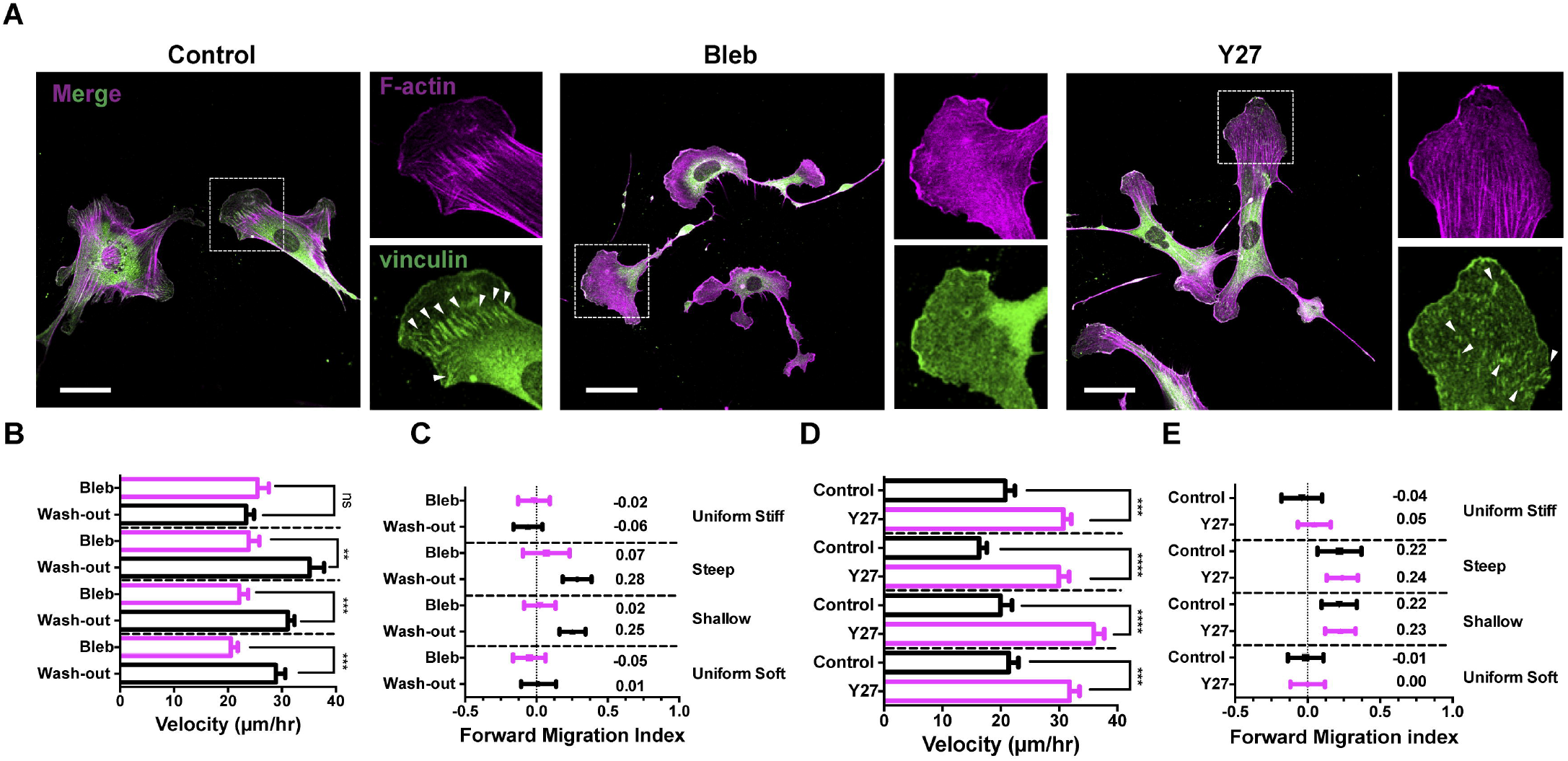
NMII activity, not ROCK, is essential for durotaxis. **(A)** Representative confocal images of JR20s in steep stiffness gradient that were treated with either DMSO (control), Bleb (30 μM) or Y27 (10 μM), fixed and stained with phalloidin (magenta) and anti-vinculin antibody (green). The area within the dashed white box is magnified. Scale bar equals 50 μm in the merged image and 25 μm in the zoom-in panels. White arrowheads mark cortical vinculin-positive FAs at the leading edge of the cells. **(B-C)** Quantification of the velocity (B) and FMI (C) of JR20s that were initially treated with Bleb (30 μM) and monitored for 18 h (Bleb) before the replacing the media without Bleb and monitored for an additional minimum 18 h-20 h (Washout) in various regions of the hydrogel is plotted. Data represents three independent experiments with ‘n’ representing individual cell value in each region - uniform stiff (n=77 for Bleb & n=94 for Washout), steep (n=40 for Bleb & n=63 for Washout), shallow (n=74 for Bleb & n=94 for Washout) and uniform soft (n=71 for Bleb & n=72 for Washout) regions. **(D-E)** Quantification of the velocity (D) and FMI (E) of JR20s that were either treated with DMSO (control) or Y27 (10 μM) in various regions of the hydrogel is plotted. Data represents four independent experiments with ‘n’ representing an individual cell’s value in each region - uniform stiff (n=50 for Control & n=83 for Y27), steep (n=53 for Control & n=94 for Y27), shallow (n=45 for Control & n=96 for Y27) and uniform soft (n=47 for Control & n=75 for Y27) regions. FMI (C & E) is plotted as values ± 95% confidence interval. Velocity (B & D) is presented as mean with s.e.m. One-way ANOVA with Sidak multiple comparison test was performed to determine statistical significance for velocity.

We next tested whether durotaxis required Rho-kinase (ROCK) activity. ROCK, a known activator of NMII (Burridge and Chrzanowska-Wodnicka, 1996; Kimura et al., 1996; Amano et al., 1996; Ridley, 2011; Burridge and Guilluy, 2016), has been implicated in durotactic response using the substrate deformation assay (Plotnikov et al., 2012). To inhibit Rho-kinase activity, we treated cells with the ROCK inhibitor Y-27632 (hereafter abbreviated Y27) seeded on hydrogels. Y27 treatment of cells led to thinner F-actin bundles and smaller FAs, but unlike Bleb, did not result in their complete disappearance (**Fig. 3A**). Interestingly, while treatment with Y27 enhanced velocity of migrating cells across all the regions, it had little impact on the FMI of cells migrating on stiffness gradients (**Figs. 3D-E**), suggesting that ROCK is dispensable for durotaxis in our system. To confirm this observation, we treated cells with an unrelated ROCK inhibitor, Rho Kinase Inhibitor III (Rockout), and found no effect on the FMI of the treated cells (**Fig. S1F**), consistent with our observations with Y27 treatment. Together, our data reveal that ROCK-based activation of NMII is dispensable for the durotaxis response of fibroblasts.

### Loss of zyxin abolishes durotactic migration

We next sought to test the role of mechano-sensitive proteins in durotaxis. FAs serve as a ‘mechanical clutch’ that connect the cell to the substrate for forward migration (Smilenov et al., 1999). Previous reports implicate FA proteins such as vinculin and paxillin in sensing substrate stiffness (Plotnikov et al., 2012). However, zyxin, a LIM-domain containing protein, is known to localize to stress fibers as well as FAs positioned along the sites of strained cell membrane (Yoshigi et al., 2005; Hoffman et al., 2012; Colombelli et al., 2009). More recently, zyxin was shown to be recruited to tensed F-actin structures via its LIM domain to regulate the elasticity of actin stress fibers in response to physical strain (Oakes et al., 2017; Uemura et al., 2011; Sun et al., 2020). We therefore hypothesized that cells use zyxin to sense stiffness gradients across the length of cell to reorganize their cytoskeleton for a directed migration response. We first stained for zyxin in cells plated on stiffness gradients, with and without Bleb treatment. In control cells, zyxin localized to both vinculin-positive FAs (white arrowheads) as well as along F-actin stress fibers (orange arrowheads) **(Fig. 4A**). As expected, upon Bleb treatment, zyxin localization to FAs and stress fibers was abolished (**Fig. 4A**), confirming a role for these structures in the localization of zyxin within fibroblasts.

**Figure 4.**
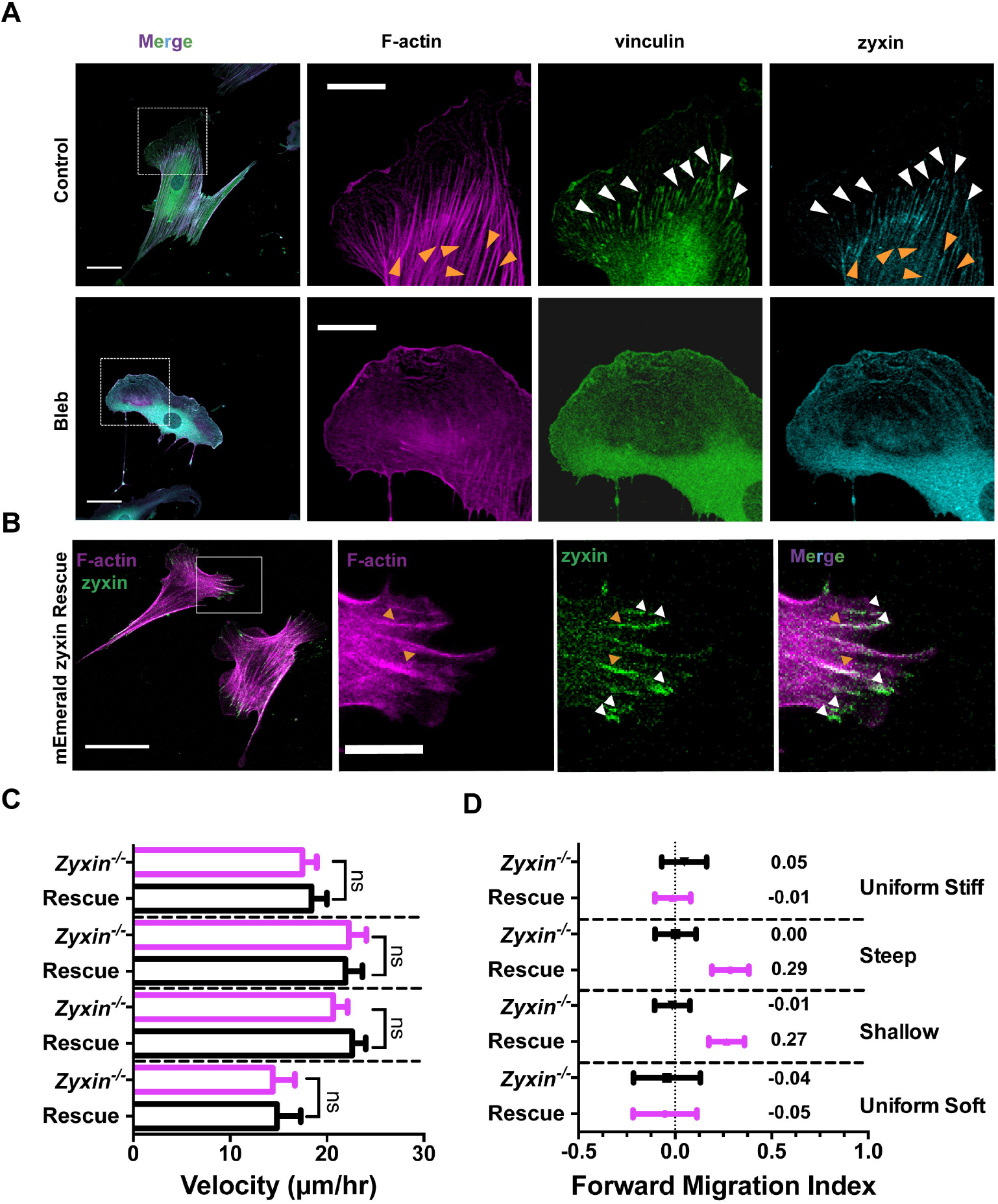
Zyxin is essential for durotaxis on different stiffness gradients. **(A)** Representative confocal images of JR20s on steep stiffness gradients that were either treated with DMSO (control) or Bleb (30 μM), fixed and stained with phalloidin (magenta) anti-vinculin (green) and anti-zyxin (cyan) antibodies. Scale bar is 50 μm for the merged image and 25 μm for zoom-in images. Dashed white box in the merge represents the region that was enlarged in magnified images. White and orange arrowheads mark the localization of zyxin along vinculin-positive FAs and actin stress fibres respectively. **(B)** Representative confocal images of mEmerald-zyxin rescue line in MEFs on steep stiffness gradient, which were fixed and stained with phalloidin. White and orange arrowheads mark the localization of zyxin to FAs and actin stress fibers respectively. Scale bar is 50 μm for the merged image and 15 μm for zoom-in images. **(C-D)** Quantification of the velocity (C) and FMI (D) of either *Zyxin^-/-^* or mEmerald-zyxin rescue (Rescue) MEF lines in various regions of the hydrogel is plotted. Data represents three independent experiments with ‘n’ representing individual cell value in each region - uniform stiff (n=48 *Zyxin^-/-^*, n=45 Rescue), steep (n=56 *Zyxin^-/-^*, n=56 Rescue), shallow (n=53 *Zyxin^-/-^*, n=79 Rescue) and uniform soft (n=24 *Zyxin^-/-^*, n=31 Rescue) regions. FMI is plotted as values ± 95% confidence interval. Velocity is plotted as mean with s.e.m and one-way ANOVA with Sidak multiple comparison test was used to determine statistical significance.

To address the functional role of zyxin in durotaxis, we used two complementary loss-of-function approaches. First, we used zyxin null (*Zyxin^-/-^*) mouse embryonic fibroblasts (MEFs) and a derivative line stably rescued with mEmerald-zyxin (**Fig. S2A**). Second, we also generated a CRISPR-based zyxin knockout (*Zyxin^KO^*) in the JR20 line, along with a *Scrambled* (*Scr*) gRNA control (**Fig. S2B**). In the rescued MEFs, mEmerald-zyxin localized to the FAs (white arrowheads) as well as to the stress fibers (orange arrowheads) (**Fig. 4B**). We analyzed the migration velocity and directionality of these cells in various regions of hydrogel system. While no difference in velocity across the regions was detected, zyxin null fibroblasts showed no durotactic response on either stiffness gradient, while the rescued derivatives showed robust durotaxis on both shallow and steep gradients (**Figs. 4C-D)**. Consistent with the *Zyxin^-/-^* MEFs, CRISPR-mediated knockout of zyxin in JR20s led to complete loss of durotaxis on both stiffness gradients, while the *Scr* control cells showed robust durotaxis (**Fig. S2C**). Therefore, zyxin is critical for a durotaxis response, irrespective of the slope of stiffness gradients.

### The role of leading-edge actin-based protrusions in durotaxis

Several previous studies have implicated leading edge actin-based protrusions, such as lamellipodia and filopodia, in sensing substrate stiffness (Oakes et al., 2018; Wong et al., 2014). To test the role of these structures in durotaxis, we investigated the functions of the Arp2/3 complex and fascin with our gradient hydrogel system. The Arp2/3 complex is the central component of branched actin networks (Goley and Welch, 2006; Rotty et al., 2013) and is critical for sensing gradients of extracellular matrix proteins (haptotaxis) (Wu et al., 2012). Conflicting data exist on the role of the Arp2/3 complex in durotaxis. Inhibition of Arp2/3 activation in cancer cells leads to impaired single cell durotaxis, while no effect on durotaxis was observed in human retinal pigment epithelial cells (hRPE) treated with an Arp2/3 inhibitor (DuChez et al., 2019; Rong et al., 2021). Fascin, an F-actin binding protein that bundles actin filaments, on the other hand, plays a critical role in the formation and protrusion of filopodia (Vignjevic et al., 2006; Jayo and Parsons, 2010). In addition, fascin can also negatively regulate NMII activity and thereby promote collective cell migration (Elkhatib et al., 2014; Lamb et al., 2021).

To functionally address the role of the Arp2/3 complex and fascin in durotaxis, we used a combinatorial genetic approach. The JR20 cell line is derived from an *Arpc2* conditional knockout mouse and expresses a tamoxifen inducible Cre recombinase (CreER^T2^) from the *Rosa26* locus. Upon tamoxifen treatment, a critical exon of the Arpc2 gene is deleted by Cre-LoxP recombination and the subsequent loss of the Arpc2 subunit leads to degradation of the entire complex (**Fig. S3A**) (Rotty et al., 2017). We used the JR20 cell line to generate a fascin knockout using CRISPR-based gene editing (*Fascin^KO^;* **Fig. S3A**). This approach allowed us to address the role the Arp2/3 complex (*Arpc2^-/-^*) and fascin (*Fascin^KO^*) in durotaxis using the same cell line and enabled us to simultaneously test the combined contributions of branched and bundled actin structures (via *Fascin^KO^Arpc2^-/-^*) to durotactic response. Surprisingly, while *Arpc2^-/-^* and *Fascin^KO^* JR20s migrated directionally in both the steep and shallow gradients, *Fascin^KO^Arpc2^-/-^* double knockout cells could durotax on the steep gradient, but not the shallow gradient (**Fig. 5A**). Further, we observed that the velocity of *Arpc2^-/-^* cells was not drastically altered, however *Fascin^KO^Arpc2^-/-^* double knockout cells displayed markedly reduced velocity on both the stiffness gradients (**Fig. 5B**), suggesting that fascin-based actin bundling is required to maintain velocity in the absence of Arp2/3 in both stiffness gradients. Interestingly, loss of fascin alone in JR20s resulted in enhanced velocity in both the stiffness gradients (**Fig. 5B**). These data establish that cells can durotax efficiently without either the Arp2/3 complex or fascin, however, the combined loss of both prevented cells from migrating directionally, specifically on shallow stiffness gradients.

**Figure 5.**
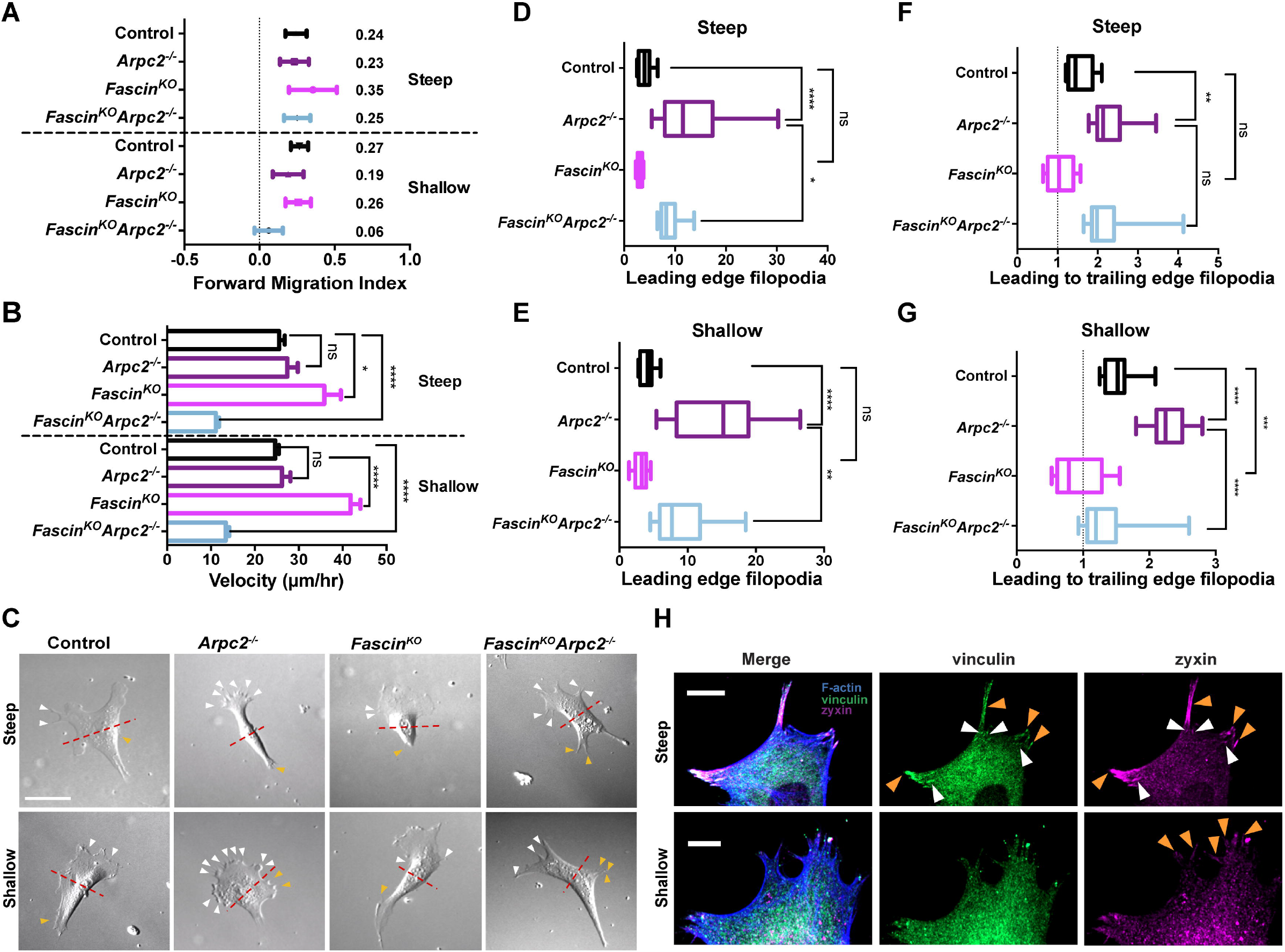
Arp2/3 complex and fascin are dispensable for durotaxis, but combined loss of both block durotaxis on shallow gradients. **(A-B)** Quantification of the FMI (A) and velocity (B) of the indicated JR20 lines in the steep and shallow gradient regions is plotted. Data represents four independent experiments for *Arpc2^-/-^, Fascin^KO^* and *Fascin^KO^Arpc2^-/-^* JR20s, whereas control JR20s are represented from twelve independent experiments. In each plot, ‘n’ represents individual cell values in each region - steep (n=159 control, n=40 *Arpc2^-/-^*, n=26 *Fascin^KO^* and n=76 *Fascin^KO^Arpc2^-/-^*) and shallow (n=208 control, n=56 *Arpc2^-/-^*, n=84 *Fascin^KO^* and n=74 *Fascin^KO^Arpc2^-/-^*) gradients. FMI is plotted as values ± 95% confidence interval. Velocity is plotted as mean with s.e.m and one-way ANOVA with Tukey’s multiple comparison test was used to determine statistical significance. **(C)** Representative DIC images of control, *Arpc2^-/-^, Fascin^KO^* and *Fascin^KO^Arpc2^-/-^* JR20 cells in steep (top row) and shallow (bottom row) gradient regions. Dashed red line separates the upper half (containing the leading edge) from the lower half (containing the trailing edge) of migrating cells. White and orange arrowheads mark the filopodia at leading and trailing edges of the cell, respectively. Scale bar equals 40 μm. **(D-E)** The number of filopodia observed at the leading edge of indicated individual JR20 lines in steep (D) and shallow (E) gradients are plotted. Pooled data from 12 independent cells collected from three independent experiments (4 cells per line per experiment for each region) **(F-G)** Ratio of leading to trailing edge filopodia of indicated JR20 individual JR20 lines in steep (F) and shallow (G) gradients. Values plotted as boxplots (D-G) with error bars of 5-95^th^ percentiles and oneway ANOVA with Tukey’s multiple comparison test was used to determine statistical significance. **(H)** Representative confocal images of *Fascin^KO^Arpc2^-/-^* double knockout JR20 cells experiencing steep (top row) and shallow stiffness gradient (bottom row), which were fixed and stained with phalloidin (blue), anti-vinculin (green) and anti-zyxin (magenta) antibodies. Images are zoom-in areas depicted from panels in Figs. S3J-K. Scale bar equals 25 μm. White and orange arrowheads point to cortical zyxin-positive FAs and zyxin-enriched filopodia observed at the leading edge of cells respectively.

To further explore the mechanism(s) behind these observations, we also analyzed the leading-edge protrusion morphology of the JR20 lines – control, *Arpc2^-/-^, Fascin^KO^* and *Fascin^KO^ Arpc2^-/-^* – migrating on both steep and shallow gradients. While the control JR20s exhibited mostly lamellipodia-based leading edge protrusions, we did observe occasional filopodia that extended along the sides of the lamellipodia as well as at the trailing edge of the cells, in both the stiffness gradients (**Fig. 5C**). In contrast, *Arpc2^-/-^* JR20s lacked lamellipodial protrusions at the leading edge and instead exhibited numerous filopodial extensions both at the leading edge as well as at the trailing edge of cells, consistent with previously published work (Wu et al., 2012; Rotty et al., 2017)(**Fig. 5C**). Using individual frames from live imaging experiments, we quantified filopodial numbers. We observed that indeed *Arpc2^-/-^* JR20s made significantly more (>5x) filopodial protrusions when compared to the control JR20s on both steep and shallow gradients (**Figs. 5C-E and S3B-E**). Lack of fascin alone had negligible impact on the filopodia number on either stiffness gradients and had similar morphology to the control JR20s (**Figs. 5C-E and S3B-E**). However, we observed that the *Fascin^KO^Arpc2^-/-^* double knockout JR20s had strikingly lower numbers of filopodia when compared to the *Arpc2^-/-^* alone controls on either stiffness gradients (**Figs. 5C-E and S3B-E**). This indicates that fascin contributes to filopodia formation and that this effect is most obvious in the *Arpc2^-/-^* background in JR20s where these structures are the dominant form of actin-based leading-edge protrusion.

Directed cell migration relies on the ability of cells to polarize migration-relevant structures relative to the external gradient cue. To test the role of the polarized distribution of filopodia in durotaxis, we compared the ratio of filopodia at the leading edge to that in the trailing edge across these cell lines. In JR20 control cells as well as *Fascin^KO^* cells, we found that leading/trailing filopodia ratio to be between ~1 to 1.5 on both the stiffness gradients (**Figs. 5F-G**), indicating that the filopodia were not polarized during the durotaxis in these two lines. Lack of the Arp2/3 complex, however, led to an asymmetric distribution of filopodia numbers in the cells with more filopodia in the leading edge of the cell than that in the trailing edge, which manifested in a much higher leading to trailing edge filopodia ratio (two or more) on both the stiffness gradients (**Figs. 5F-G**). Importantly, the asymmetric filopodial distribution was attenuated in *Fascin^KO^Arpc2^-/-^* double knockout JR20s when plated on the shallow gradient, but not on the steep gradient (**Figs. 5F-G**). We also addressed whether filopodial length rather than filopodial number may be polarized during durotaxis. In control and *Fascin^KO^* cells, we found no significant difference in the length of filopodia at the leading or trailing edge either shallow or steep gradients (**Figs. S3F and S3H-I**). However, on shallow gradients, lack of Arp2/3 resulted in a modest yet significant increase in the filopodia length at the leading edge, that was largely independent of fascin (**Fig. S3G**), pointing to other potential regulators of filopodia at play here. Importantly, on the shallow gradient, fascin is required for the maintenance of asymmetric filopodia distribution in the absence of branched actin networks.

Since leading edge protrusion is linked to the formation of cell-matrix adhesions via FAs (Schwarz and Gardel, 2012), we stained for FAs and zyxin in conjunction with F-actin in cells lines lacking Arp2/3, fascin, or both, along with controls on both the stiffness gradients. On the steep gradient, we observed zyxin localizing to vinculin-positive FAs at the leading edge (**Fig. S3J**). Furthermore, we also observed zyxin localize to filopodial shafts, specifically in *Arpc2^-/-^* and *Fascin^KO^Arpc2^-/-^* double knockout lines, on both the stiffness gradients **(Figs. 5H and S3J-K**). Importantly, *Fascin^KO^Arpc2^-/-^* double knockout cells specifically lacked zyxin-positive FAs only on the shallow, but not steep, stiffness gradient (**Figs. 5H and S3J-K**), suggesting a potential role for fascin in the formation of cortical zyxin-positive FAs to promote durotaxis, specifically in the shallow gradient.

The work presented in this study provides several technical and conceptual advances for the field of mechanically directed cell migration. Our new photopolymerized hydrogel system is robust and simple to implement in any cell biology lab. This should enable more labs to interrogate molecular mechanism(s) of durotaxis across a variety of cell types. A key feature of this system is the ability to generate two distinct gradients in one gel, along with uniform soft and stiff regions to serve as internal controls. As typified by our *Fascin^KO^Arpc2^-/-^* results, some perturbations will affect cells durotaxing on the shallow gradients, but not the steep gradients. The uniform areas of the gel allow the appropriate controls to ensure that directed migration can be detected with confidence, as well as allowing cell properties that vary with stiffness to be measured within a single dish of cells. Our experimental set-up is also amenable to high resolution fixed and live-cell imaging, allowing molecular scale information in durotaxing cells to be obtained. Finally, a major strength of this system is the ability to tune the absolute stiffness and gradient properties by simply varying the timing of illumination of the polymerizing gel. This will allow investigators to tune the hydrogel properties to address their specific questions and accommodate new cell types that prefer different stiffness ranges.

From a conceptual point of view, our data support several important conclusions and prompt new questions. Our data reaffirms the essential role of NMII in durotactic response but raises important questions about the regulation of this activity. Using two distinct inhibitors of Rho-kinase, we conclude that ROCK-mediated regulation of NMII is not required for single cell fibroblast durotaxis. This is surprising since this regulatory input is thought to be particularly important for connecting RhoA-based signaling to cell contractility (Riento and Ridley, 2003). Perhaps other regulatory inputs such as myosin light chain kinase (MLCK) might be critical for NMII activity during durotaxis. Alternatively, cells may simply utilize basal NMII activity to actively probe the mechanical landscape of the substrate, but do not require spatial or temporal regulation of NMII activity to durotax. Our data also support the role of the mechano-sensitive protein zyxin in the process of durotaxis (Yip et al., 2021). Taken together with previous studies of focal adhesion proteins vinculin and paxillin in the process of durotaxis, our data are consistent with the notion that cells use focal adhesion structures to sense the local stiffness of the substrate. In future studies, it will be important to spatially and temporally resolve the role of focal adhesions versus the role of adjacent contractile actin bundles where zyxin also localizes in the process of durotactic sensing. Our new durotaxis assay should facilitate the enumeration of all the proteins required in these structures to execute durotactic migration. A broader question about this process is how sensing events that occur locally via focal adhesions translates to whole cell polarity and directed migration specifically on stiffness gradients (SenGupta et al., 2021).

Our study also speaks to the role of actin-based protrusions at the leading edge in the process of durotaxis. Of particular importance is the finding that the Arp2/3 complex is dispensable for fibroblast durotaxis. This stands in stark contrast to extracellular matrix (ECM) haptotaxis where the Arp2/3 complex is absolutely required for response to these directional migration cues (Wu et al., 2012; King et al., 2016). In physiological situations, durotaxis and haptotaxis are likely to be highly intertwined as the content/amount of ECM and its mechanical properties are interdependent. Here, we have deconvolved these processes and revealed the striking diversity of possible mechanisms of directed migration. Our *Arpc2^-/-^* JR20s completely lack lamellipodia and instead employ a filopodia-driven protrusion system, and yet are capable of efficient durotaxis on steep and shallow stiffness gradients. This implies that either type of leading edge protrusion can support durotactic migration. Consistent with this idea, *Fascin^KO^* cells are also capable of efficient durotaxis on both steep and shallow gradients. Interestingly, only when both components are deleted (*Fascin^KO^Arpc2^-/-^*) do defects begin to become apparent. We postulate that the shallow gradient in our system presents a more difficult directed migration challenge for cells and that properly regulated leading edge protrusions (either lamellipodia or filopodia) are required for efficient durotaxis in this situation. The known interplay between leading edge protrusions and the formation of focal adhesions may be key to abnormal protrusions in the *Fascin^KO^Arpc2^-/-^* cells leading to inefficient focal adhesion formation just behind the leading edge. Together, the work presented here provides critical insights into the mechanism of durotaxis and provides an experimental approach that lays the foundation for future studies of this process.

## MATERIALS AND METHODS

### Reagents and materials

Antibodies for western blotting and immunofluorescence: anti-GAPDH (AM4300), AlexaFluor 488-conjugated anti-mouse (A11001), Rhodamine Phalloidin (R415), and AlexaFluor 405 Phalloidin (A30104) were from ThermoFisher Scientific; anti-ARPC2 (07-227) and anti-Fascin (55K2) antibodies were from Millipore (MAB3582); anti-Vinculin (hVIN-1) antibody was from Sigma-Aldrich; Cy5-conjugated anti-rabbit (A10523) antibody was from Jackson Laboratory; IRDye 800CW-conjugated anti-Rabbit (92532211) and IRDye 680RD-conjugated anti-Mouse (92568070) antibodies were from LI-COR; and anti-Zyxin (Polyclonal rabbit serum B71) was a gift from Mary Beckerle (Hoffman et al., 2003). Human fibronectin (356008) was purchased from Corning. Cy5-labelling kit purchased from GE Healthcare. Para-aminoblebbistatin (ax-494682) and Rho-kinase III inhibitor (17922) were purchased from Axol Bioscience and Cayman Chemicals, respectively. 4-Hydroxytamoxifen (H6278-10MG), Y-27632 (Y0503) and DMSO (D2650) were purchased from Sigma Aldrich. Lithium Phenyl(2,4,6-trimethylbenzoyl) phosphinate (LAP; L0290) was purchased from TCI chemicals. Sulfosuccinimidyl 6-(4’-azido-2’-nitrophenylamino) hexanoate (Sulfo-SANPAH; 22589) from ThermoFisher Scientific. Acrylamide solution (40%) and bis-acrylamide solution (2%) from Fisher Scientific. 30 mm coverslips were obtained from Bioptechs (30-1313-0319) and 500μm^2^ gridded coverslips (10816) from ibidi. 12- and 18-mm coverslips, 3-Aminopropyl triethoxysilane (APTES; 98% solution. A3648), Gluteraldehyde (50% Aqueous solution. 16320), and 20% Paraformaldehyde were purchased from Electron Microscopy Sciences. Triangular silicon nitride atomic force microscope cantilevers with 4.5 μm polystyrene particle and 0.06 N/m spring constant were purchased from NovaScan.

### Cell Culture

All cells were cultured and imaged in high-glucose DMEM (4.5 g/L D-glucose, 0.584 g/L L-glutamate, 110 mg/L sodium pyruvate; Gibco) supplemented with 10% fetal bovine serum (MedSupply Partners), 100U/ml Penicillin/Streptomycin (Gibco), and 1% Glutamax (Gibco) at 37° C and 5% CO_2_. Cells were split using 0.25% trypsin-EDTA (Gibco).

### Generation of CRISPR Cas 9-mediated gene knockouts

The following guide RNA sequences were used to target indicated genes: 5’-GCTACGCGCATCTGAGCGCG-3’ for *Fascin*, 5’CTGCGGGGCGTAAAACGCGG3’ for *zyxin*, and 5’GCGCTCTACCGATACCGCTC3’ used as a non-targeting (*Scr*) control. gRNA targeting *Fascin* was cloned into pLentiCRISPRv2 (Addgene # 52961; a gift from Feng Zhang) and the gRNAs for *Zyxin* or *Scrambled* were cloned into pTLCV2 (Addgene # 87360; a gift from Adam Karpf).

### Cloning, transfection, and lentiviral infection

The following plasmids were used to generate zyxin rescues:

mEmerald-Zyxin-C-14 construct (Addgene # 54318; a gift from Michael Davidson) was cloned into the lentiviral vector pLL5.0 (Vitriol et al., 2007) via Gibson assembly. Rescue constructs or CRISPR constructs were transfected alongside pCMV-V-SVG, pRSV-REV, and pMDLg/pRRE (500 ng each) into HEK293Ts using X-tremegene HP Transfection reagent (Roche). Lentivirus was harvested 72 hours later and subsequently used to infect JR20s supplemented with 4 μg/ml Polybrene. For the generation of *Fascin^KO^*, JR20s expressing CRISPR components targeting Fascin were selected using 2 μg/ml puromycin treatment for 48 hours before being subjected to clonal selection and expansion. For the generation of the *Scr* and *Zyxin^KO^*, infected JR20s were subject to two rounds of puromycin selections (2.5 μg/ml and 5 μg/ml) followed by three rounds of doxycline induction at 10, 25, and 50 μg/ml. GFP-positive cells were FACs sorted at the end of each round of doxycycline induction and propagated for targeted assays.

### Tamoxifen-induced generation of Arpc2^-/-^ and Fascin^KO^Arpc2^-/-^ lines

JR20s were split into two 6 cm dishes (50,000 cells each). One dish was treated with 2 μM of 4-hydroxy-tamoxifen (4-OHT), and the second dish was treated with ethanol (0.0002% v/v; control). After 48 h, the media was replaced with a second dose of 4-OHT or ethanol in either the treated or control dish, respectively. 48 h post second dosing, cells were harvested for targeted assays. To prevent inadvertent compensatory mechanisms from occurring, *Arpc2^-/-^* cells were only used up to two weeks post 4OHT treatment. *Fascin^KO^* cell line was subjected to the same protocol to generate *Fascin^KO^Arpc2^-/-^* line.

### Western Blotting

Cells (2.5×10^5^) were plated in 6cm dishes, incubated overnight, washed with 1X DPBS the next day, and subsequently scraped with ice cold RIPA buffer (50 mM Tris pH 8, 150 mM NaCl, 0.5% deoxycholate, 0.1% SDS, 1% NP-40; all chemicals from Sigma Aldrich) supplemented with protease and phosphatase inhibitors (Roche). Lysates were left on ice for 15 minutes before addition of Laemmli buffer and boiled for 10 minutes. Samples were run on 4-12% gradient SDS-PAGE gels (BIO-RAD), transferred to nitrocellulose membranes (BIO-RAD) and blocked with non-fat dry milk (LabScientific; 5% in Dulbecco’s phosphate buffered saline or DPBS) for 1h at room temperature (RT) before overnight incubation at 4°C with primary antibodies (diluted in 1% bovine serum albumin containing PBS supplemented with 0.1% Tween-20, v/v (PBST), and 0.01% sodium azide). Blots were washed with PBST and then incubated at RT with fluorescent-conjugated secondary antibodies for 1 h. Blots were then visualized using LI-COR Odyssey system.

### Engineered photoactivated hydrogel system

#### Instrument set up

An Olympus collimator (C2000Z-ADP/IX) was attached to and beneath a Thorlabs LED UV light source (365 nm) (see **Fig. S1A**), which was placed 3.5 cm above the sample. The Thorlabs LED power supply potentiometer was set at 0.35 Amp and the main rheostat turned to 50%. Using a light meter, we measured that 0.2 mW of light was being illuminated over a circular area of 2cm^2^. From this, we determined the optimum light flux delivered to the sample as 0.1 mW/cm^2^.

#### Glass activation

30 mm coverslips were washed with 100% ethanol, air dried, and air plasma cleaned for 2-3 mins. Coverslips were immersed in 0.5% APTES (20mins) followed by washing with deionized (DI) water (2 times, 5 min each). Coverslips were immersed in 0.5% Glutaraldehyde (45 mins) and washed afterwards (immersed in water 3 times, 10 min each, and air dried). Activated coverslips were stored in 4 degrees.

#### LAP acrylamide mixture

A 12% acrylamide and 0.6% bis-acrylamide solution in DPBS was prepared. LAP solution (0.5% v/v) was made using the acrylamide mix in a glass vial (500 μL volume). The glass vial containing the mixture was covered in aluminum foil, protected from light exposure, and de-gassed in vacuum chamber for 10-15 min.

#### Substrate preparation

The chemically activated 30 mm glass coverslip was placed in a 3D printed sample holding mold (see dimensions in **Fig. 1A**) and a 15 μL drop of the LAP acrylamide mixture was placed in the center of the coverslip. A 12- or 18-mm glass coverslip was gently lowered over the drop to create a sandwich. The 3D-printed mold with a house-shaped void on the top surface was placed with the ‘house roof’ pointing left and inserted into the path of light source. The bottom half of the house-shaped void was covered with an opaque paper mask. The sandwich was exposed for 3 mins. At the 3-minute mark, the paper slip was quickly and completely removed, and sample was further exposed to light for another 45 seconds. Light was quickly turned off, the top coverslip (12- or 18-mm) was gently dislodged with fine forceps, and the gel was rinsed and stored in DPBS at 4°C until further use. STL files of 3D printed parts are available upon request.

#### Fibronectin crosslinking

Stored gels were gently washed thrice with DI water. 1 mg/mL Sulfo-SANPAH solution was prepared in DI water from a concentrated 25 mg/mL stock in DMSO stored at −80°C. The Sulfo-SANPAH solution was centrifuged at maximum speed (15,000 RPM) and passed through an 0.20 or 0.45 μm filter to remove particulate material. Filtered Sulfo-SANPAH solution was added to the gel surface and illuminated in a UV Stratalinker (Stratagene) oven for 25-30 minutes. Gel was then quickly transferred from the oven to tissue culture hood, rinsed 3x with sterile water and then incubated in 10 μg/mL human fibronectin (FN) made in DPBS for 90 min at 37°C. Remaining fibronectin solution was then aspirated, gel was rinsed twice with DPBS and left immersed in DMEM prior to seeding and imaging of cells.

### Analyses of gel elastic modulus

#### Atomic force microscopy approach

Force measurements were performed with an Asylum Research MFP (3D) instrument. A triangular silicon nitride cantilever with a 4.5 μm polystyrene particle and 0.06 N/m spring constant was used. Before collecting measurements, the cantilever sensitivity was calibrated by measuring the force–distance slope in the AFM software on a clean glass slide with a drop of (1x) DPBS. For measurements of the gradient, the house-shaped gel was covered with a drop of DPBS to reduce random adhesion of tip to the gel and thereby improve force measurements. For each position, 3 force curves were acquired with a trigger force of 10 nN, an approach speed of 1 μm/s and a maximum set point of 0.5 or 1.0 V. The Hertz contact model was used to extrapolate the elastic moduli from the force-distance measurements on the Igor Pro software (version 6.22).

#### Nanoindentation approach

Hydrogels were generated on 500 μm^2^ gridded coverslips (Ibidi) as described previously for regular glass coverglass and was submerged in DPBS. The location of the “Line” on the gel with respect to gridded coverglass was identified via brightfield imaging using a 10x air objective (0.45; Plan-Apochromat). The identified region was marked and subsequently used to orient the gridded coverslip for nanoindentation measurements. Nanoindentation was performed with an Optics11 Piuma Nanoindenter with a 9.5 μm radius spherical tip and a spring constant of 0.30 N/m. The Piuma Nanoindenter was first calibrated against a glass petri dish. The gradient gel on the gridded coverglass was then aligned on the nanoindenter such that the tip of the nanoindenter was directly above the marked ‘line of altered contrasts’ region on the coverglass (spanning 5 grid squares), using an inbuilt USB camera. Nanoindentation measurements were acquired every 50 microns, moving vertically from top to bottom of the house-shaped gel (spanning 10 grid squares: ~5mm). Images were also acquired at the first and last contact point of the probe for later alignment of the elastic modulus data and grid lines. The Hertzian contact model was used to compute elastic modulus from the reported force indentation measurements on the Dataviewer software V2.3.

### Cell migration analysis on hydrogels

Hydrogels were coated with 10 μg/ml uniform human FN (Corning) for 90min at 37°C and rinsed twice with DPBS before seeding cells. Cells were sparsely seeded and allowed to spread for 2-3 h on gels, before being imaged at 37°C and 5% CO_2_ using a 20x objective on an Olympus VivaView FL microscope for a period of 18-30 h with 10 min interval between each frame. Single cells from each field of view (FOV) were tracked manually using the *Manual Tracking* plugin in ImageJ. Cell tracking was aborted whenever cells collided with each other or migrated outside the field or divided or died during acquisition in each FOV. Also, cells adhered to each other were not tracked. Velocity, forward migration index (FMI) and persistence (D/T) were extracted from raw tracking data using the *Chemotaxis Tool* plugin in ImageJ.

### Preparation of Cy5-conjugated Fibronectin

Cy5-labelling kit purchased from GE Healthcare was used for conjugation of human fibronectin (Corning), as previously described (Wu et al., 2012).

### Immunofluorescence, confocal microscopy, and image analysis

Hydrogels were coated with Cy5-conjugated FN for 90 minutes at 37°C and images were tiled to acquire the entire house-shaped gel using a 5x air objective (0.16; Plan-Apochromat name) on Zeiss Airyscan 880 confocal microscope. For F-actin and immunofluorescence staining, cells were plated at approximately 40% confluency and allowed to spread overnight. Cells were then fixed at 37°C for 10 minutes with paraformaldehyde (4% final concentration in the growth media). Cells were then permeabilized with 0.02% Triton-X 100 (Fisher Scientific) at 37°C for 10 minutes and blocked with DPBS containing 5% BSA and 5% normal goat serum (Gibco), for 30 mins at 37°C. Primary antibodies were diluted in 1% BSA in DPBS and added to the gel and incubated for ~2 h at 37°C. After rinsing the gel thrice using DPBS, DPBS containing 1% BSA and manufacturer recommended dilutions of secondary antibody and rhodamine phalloidin were added to gels and incubated for 1 h at 37°C. Gels were then washed thrice and stored in DPBS at 4 degrees until imaging. Confocal microscopy was performed on the Zeiss Airyscan 880 microscope using a 40x oil objective (1.3; EC Plan-Neofluar) and a 20x objective (0.8; Plan-Apochromat). Images were processed using Imaris, MATLAB and/or Fiji softwares. For representation, images were adjusted for brightness and contrast across the whole image as well as to similar extent between conditions. Images were cropped using Fiji or PowerPoint (Microsoft 365) to be presented as magnified insets.

### Measurement of Cy5-Fibronectin uniformity across hydrogels

The integrated density profile of cy5-fibronectin was extrapolated from Fiji. Briefly, a boxed region of interest (ROI) was set at the top, center, and bottom of the hydrogel. The integrated density profiles of the three regions (top, center, and bottom) were plotted alongside each other and Tukey’s multiple comparison test was utilized to see if there are significant differences between the profiles across the three different regions of the gel.

### Circularity and Nematic Order Analysis

Circularity and Nematic order analysis (NA) were evaluated in JR20s stained with phalloidin and acquired using confocal microscope as mentioned above. For cell circularity, MATLAB image processing toolbox was used where cells were individually masked using segmentation tool and then circularity was computed from each mask using the region props where circularity was computed as (4*pi*Area)/(Perimeter^2). For NA, the local orientation of actin and coherence were determined at each pixel from the segmented image using a custom-build MATLAB program generously provided by the Ladoux lab (Doss et al., 2020), where the order parameter was S=<cos(2|θ–θavg|)>and θ avg is the average angle of actin filaments. All averages were weighted by the fluorescence intensity across the cell.

### Filopodia Analysis

JR20 cells from time-lapse DIC (Differential Interference Contrast) imaging experiments mentioned above were utilized for filopodia analysis. Each JR20 line migrating on either steep or shallow stiffness gradients were independently analyzed for filopodia. Individual cells were first split into upper half (leading edge) and bottom half (trailing edge), based on their direction of migration. Filopodia-like extensions, typically 2-10 μm in length, were manually identified, measured, and counted in the leading as well as trailing edges of a migrating cell in each line. For each cell, a minimum of four frames (~10 mins between each frame) were analyzed for filopodia numbers and distribution, and four cells per gradient region for each JR20 line were analyzed in each experiment. Membrane extensions >10 μm as well as large retraction fibers were excluded from analysis. Total filopodia represents the sum of filopodia observed at the leading and trailing edges in each cell.

### Statistical Analysis

All statistical analyses were conducted with GraphPad Prism (version 6.0). In box and whisker plots, data points represent 5^th^-95^th^ percentile range. Unless otherwise stated, statistical significance was determined using a one-way ANOVA test involving Tukey’s multiple comparison test, with p-values < 0.05, <0.001 and <0.0001 represented as *, ** and *** in the graphs.

## ACKNOWLEDGEMENTS

We thank members of the Bear lab for scientific feedback on this work. We gratefully acknowledge support from a National Institutes of Health grant to J.E.B. (R35GM130312). This work was in part performed in part at the Chapel Hill Analytical and Nanofabrication Laboratory, CHANL, a member of the North Carolina Research Triangle Nanotechnology Network, RTNN, which is supported by the National Science Foundation, Grant ECCS-2025064, as part of the National Nanotechnology Coordinated Infrastructure, NNCI. We thank Benoit Ladoux’s lab for providing the Nematic Order MATLAB code. We also thank Mary Beckerle for the gift of zyxin^-/-^ MEF cell lines.

## AUTHOR CONTRIBUTIONS

Conceptualization: R.M.H and J.E.B; Study design: R.M.H, B.C.S and J.E.B; Experimentation and analysis: R.M.H, B.C.S, M.A.H, Z.T.K., M.T.B and W.R.L; Manuscript preparation: R.M.H, B.C.S and J.E.B; Funding acquisition: J.E.B.

## SUPPLEMENTARY FIGURE LEGENDS

**Figure S1. (A)** A picture of the hardware set-up for the generation of engineered house-shaped hydrogel for the study of single cell durotaxis response. **(B)** Tiled image of the engineered house-shaped hydrogel on a 500 μm^2^ gridded coverslip. Dashed orange rectangle marks the line of altered contrast **(C-D**) A representative tiled confocal image (C) and the integrated density profile of Cy5-conjugated fibronectin (D) across three 1000 μm regions (see black lines in panel C), from two independent fibronectin-coated house-shaped hydrogels. **(E)** Quantification of the FMI of JR20s that were initially untreated (control) and monitored for 20 h before being replaced with media containing Bleb (30 μM) and monitored for an additional 20 h (Bleb wash-in) in various regions of the hydrogel is plotted. Data represents two independent experiments with ‘n’ representing individual cell value in each region - uniform stiff (n=50 control, n=52 Bleb wash-in), steep (n=63 control, n=87 Bleb wash-in), shallow (n=45 control, n=56 Bleb wash-in) and uniform soft (n=44 control, n=58 Bleb wash-in). **(F)** Quantification of the FMI of JR20s that were either treated with DMSO (control) or Rockout (50 μM) and monitored for 30 hours. Data represents four independent experiments with ‘n’ representing individual cell value in each region-uniform stiff (n=44 control, n=38 Rockout), steep (n=65 control, n=44 Rockout), shallow (n=44 control, n=49 Rockout), and uniform soft (n=39 control, n=24 Rockout). FMI values (E-F) are plotted as mean values ± 95% confidence interval.

**Figure S2. (A-B)** Western blot analysis of the amounts of zyxin in the lysate obtained from NIH3T3 fibroblasts, JR20s as well as *Zyxin^-/-^*, mEmerald-zyxin rescues and mScarlet-zyxin rescue MEFs (A) as well as in the lysates from *Scr* and *Zyxin^KO^* JR20 lines lysates (B). Gapdh served as a loading control. **(C)** Quantification of the FMI of *Scr* and Z*yxin^KO^* JR20 lines in various regions of the hydrogel is plotted. Data represents four independent experiments with ‘n’ representing individual cell value in each region - uniform stiff (n=35 *Scr*, n=41 *Zyxin^KO^*), steep (n=45 *Scr* n=42 *Zyxin^KO^*), shallow (n=49 *Scr* n=59 *Zyxin^KO^*) and uniform soft (n=44 *Scr*, n=37 *Zyxin^KO^*) regions and plotted as values ± 95% confidence interval.

**Figure S3. (A)** Western blot analysis of the amounts of Arpc2 and fascin expressed in the lysates from *Scr*, control, *Arpc2^-/-^, Fascin^KO^* and *Fascin ^KO^Arpc2 ^-/-^* double knockout JR20s. Gapdh served as a loading control. **(B-I)** Analysis of the number of filopodia at the trailing edge (B & C) as well as the total number for each cell (D & E) in the indicated individual JR20 line within steep (B & D) and shallow (C & E) gradient regions of the hydrogel, respectively. Analysis of the length of filopodia at the leading edge (F & G) and trailing edge (H & I) for each cell of the indicated individual JR20 line within steep (F & H) and shallow (G & I) gradient regions of the hydrogels, respectively. Pooled data from 12 independent cells collected from three independent experiments (4 cells per line per experiment for each region) are plotted as 5th-95^th^ percentile of mean and one-way with Tukey’s multiple comparison test was used to determine statistical significance. **(J-K)** Representative confocal images of indicated JR20 lines in steep (J) and shallow (K) stiffness gradient, which were fixed and stained with phalloidin (blue), anti-vinculin (green) and anti-zyxin (magenta) antibodies. Scale bar is 50 μm for merge channel and 25 μm for zoom-in. White and orange arrowheads point to cortical zyxin-positive FAs and zyxin-enriched filopodia observed at the leading edge of cells respectively.

