## Supplementary figures and images for "Zyxin and non-muscle myosin are required for single fibroblast durotaxis, but Rho-kinase activity and the Arp2/3 complex are dispensable"

### Supplementary Figure S1

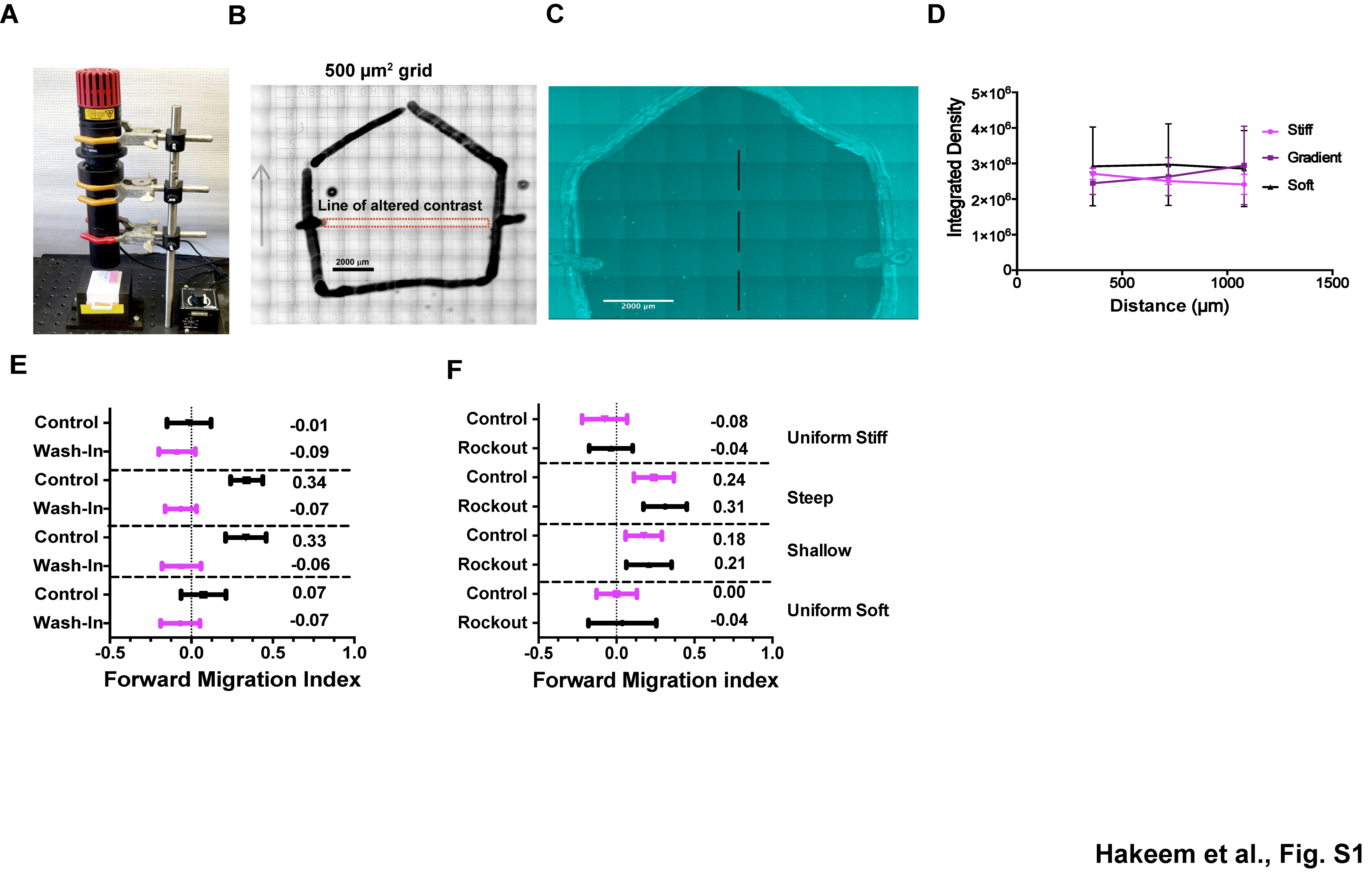

### Supplementary Figure S2

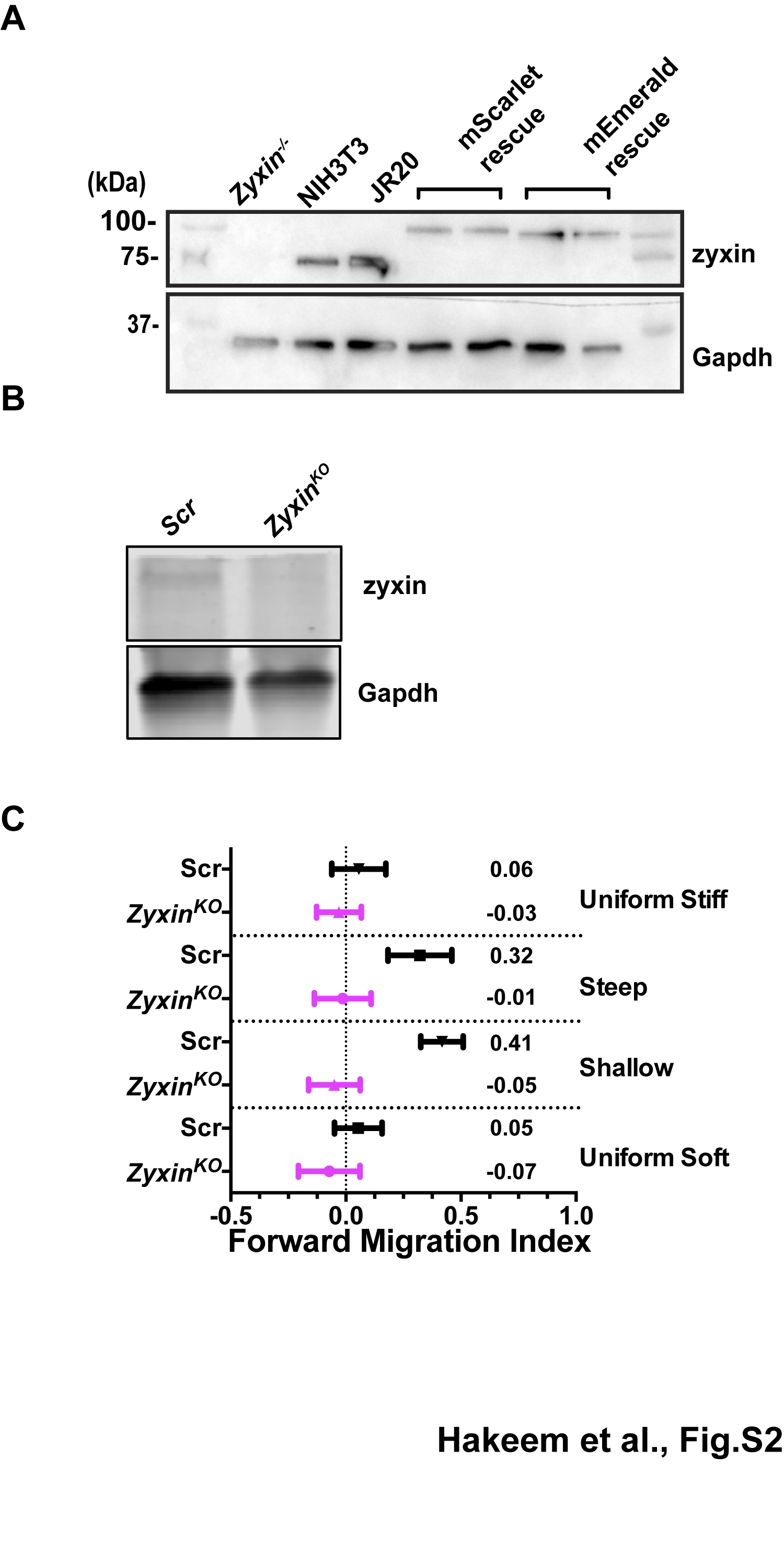

### Supplementary Figure S3

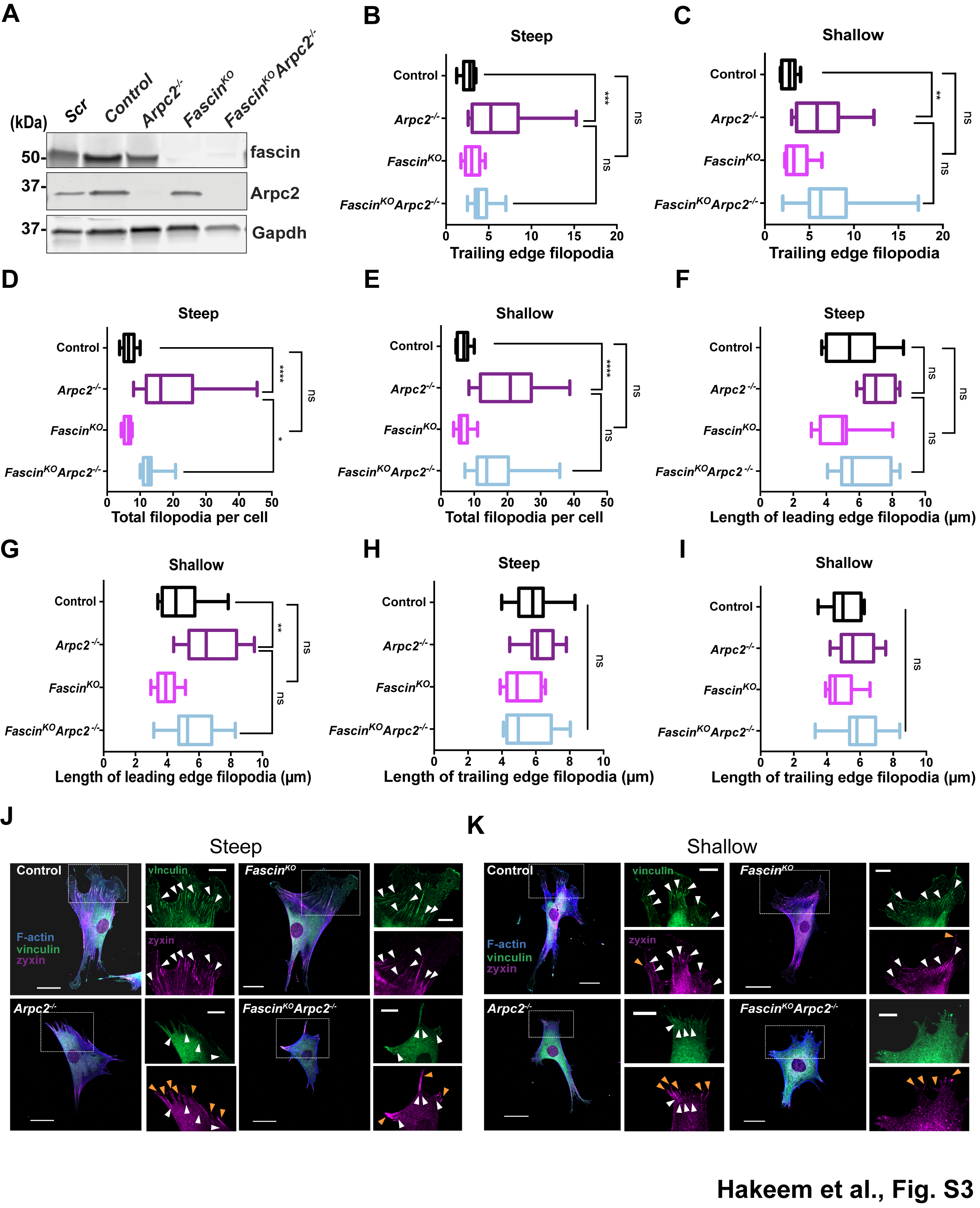
